# Epistatic genetic interactions govern morphogenesis during sexual reproduction and infection in a global human fungal pathogen

**DOI:** 10.1101/2021.12.09.472005

**Authors:** Sheng Sun, Cullen Roth, Anna Floyd Averette, Paul Magwene, Joseph Heitman

## Abstract

Cellular development is orchestrated by evolutionarily conserved signaling pathways, which are often pleiotropic and involve intra- and inter-pathway epistatic interactions that form intricate, complex regulatory networks. *Cryptococcus* species are a group of closely-related human fungal pathogens that grow as yeasts yet transition to hyphae during sexual reproduction. Additionally, during infection they can form large, polyploid titan cells that evade immunity and develop drug resistance. Multiple known signaling pathways regulate cellular development, yet how these are coordinated and interact with genetic variation is less well understood. Here, we conducted quantitative trait locus (QTL) analyses of a mapping population generated by sexual reproduction of two parents, only one of which is unisexually fertile. We observed transgressive segregation of the unisexual phenotype among progeny, as well as a novel large-cell phenotype under mating-inducing conditions. These large-cell progeny were found to produce titan cells both *in vitro* and in infected animals. Two major QTLs and corresponding quantitative trait genes (QTGs) were identified: *RIC8* (encoding a guanine-exchange factor) and *CNC06490* (encoding a putative Rho-GTPase activator), both involved in G-protein signaling. The two QTGs interact epistatically with each other and with the mating-type locus in phenotypic determination. These findings provide insights into the complex genetics of morphogenesis during unisexual reproduction and pathogenic titan cell formation and illustrate how QTL analysis can be applied to identify epistasis between genes. This study shows that phenotypic outcomes are influenced by the genetic background upon which mutations arise, implicating dynamic, complex genotype-to-phenotype landscapes in fungal pathogens and beyond.

**Significance statement:** Cellular development is regulated by a complex web of signaling pathways that respond to both intracellular and extracellular cues. Morphological transitions in pathogenic fungi, such as those observed during sexual reproduction or in response to the host environment, offer tractable models for understanding the principles that govern eukaryotic cell development and morphogenesis. Using the human fungal pathogen *Cryptococcus deneoformans* as a model and applying QTL analysis, we defined novel genes and gene-gene interactions involved in the yeast-hyphal transition and titanization, two morphological developments that are important for adaptation, pathogenesis, and evolution of this fungal pathogen. Our study provides new insights into the conservation and complexity of key signaling pathways in regulating cell development in fungi, as well as other eukaryotes.

## Introduction

Evolutionarily conserved signaling mechanisms, such as those involving G-protein coupled receptors (GPCR), cyclic AMP (cAMP), and mitogen-activated protein kinases (MAPK), govern cellular development. Changes in these signaling networks often have broad developmental effects due to the pleiotropic nature of many of these pathways, such as those involved in responses to nutrient stress in yeast and other fungi (1). Phenotypic outcomes of genetic changes are often dependent on the genetic background in which they appear, and such background effects have been identified for a variety of phenotypes such as gene essentiality, and in many different model systems including yeasts, nematodes, fruit flies, mice, and human cell lines (2-14). These background effects impede our ability to understand the phenotypic outcomes of genetic changes, which can be further complicated by the higher-level interactions that exist among modifiers. Identifying such modifiers, as well as understanding the underlying mechanisms of their function and interactions, are critical to establish the basis of phenotypic and genotypic variation.

In the fungal kingdom, sexual reproduction is a complex developmental process and occurs through two mechanisms: heterothallism and homothallism. For heterothallic species, sexual reproduction involves two individuals of different and compatible mating types, similar to sexes in animals. Heterothallic sexual reproduction brings together the genomes of two individuals, produces recombinant progeny through meiosis, and can generate genotypic and phenotypic variation, thus facilitating natural selection. On the other hand, homothallic species can initiate and successfully complete sexual reproduction from a single individual, through mechanisms like mating-type switching, fusion of compatible mating type alleles, co-segregation of compatible mating-type alleles during meiosis, co-inheritance of nuclei with compatible mating types in progeny, and unisexual reproduction (15, 16). While homothallic sexual reproduction typically produces inbred offspring that are genetically uniform and identical to the parent, unisexual reproduction can also occur between two genetically different individuals of the same mating type, producing recombinant progeny through meiosis (17-19). Morphological changes during sexual development are complex traits that can be influenced both environmentally and genetically, and typically require intricate control by several evolutionarily conserved cellular pathways. These include GPCRs, cAMP, and MAPK signaling pathways that mediate pheromone responses, sense environmental stimuli (e.g. pheromone and nutrient) and transduce these signals to drive sexual development in both homothallic and heterothallic sexual reproduction (20-22).

Pathogenic *Cryptococcus* species are basidiomycetous yeasts that comprise three currently recognized species or species complexes: *Cryptococcus neoformans* (formerly *C. neoformans* var. *grubii*, serotype A), *Cryptococcus deneoformans* (formerly *C. neoformans* var. *neoformans*, serotype D, and *Cryptococcus gattii* (formerly *C. neoformans* var. *gattii*, serotypes B and C) (23-25). These species are opportunistic pathogens of humans and other animals that cause systemic infection involving dissemination to the central nervous system to cause meningoencephalitis. Collectively, *Cryptococcus* species infect an estimated 220,000 to one million people each year, causing 140,000 to 600,000 deaths annually (26, 27). The primary infectious propagule is hypothesized to be desiccated yeasts or spores produced through sexual reproduction. Most basidiomycete species have a tetrapolar mating system in which mating type is determined by alleles at two unlinked mating type (*MAT*) loci: a *P/R* locus encoding pheromones and pheromone receptors and an *HD* locus encoding transcription factors controlling cell type identity and sexual development. In contrast, *Cryptococcus* species have a bipolar mating system that evolved from an ancestral tetrapolar mating system through chromosomal rearrangements and fusion of the *P/R* and *HD* loci. The mating type of *Cryptococcus* is determined by the presence of the **a** or α allele at the *MAT* locus. Compared to most basidiomycetes, the *Cryptococcus MAT* locus is large (>100 kb) and contains both the classic genes associated with *MAT* in tetrapolar basidiomycetes (pheromones (*MF*), pheromone receptors (*STE3*), and homeodomain transcription factors (*SXI1* and *SXI2*)), as well as additional genes that have been recruited into the *Cryptococcus MAT* during the tetrapolar to bipolar mating system transition (28-32).

In heterothallic sexual reproduction, also called bisexual reproduction or opposite-sex mating, two *Cryptococcus* cells of opposite mating type conjugate and the resulting zygote initiates sexual development, involving hyphal growth, basidia formation, and spore production. In the *C. deneoformans* complex, a form of homothallic sexual reproduction, called unisexual reproduction, has also been discovered (19). In this case, sexual reproduction can be initiated from a single cell of either **a** or α mating type. Sexual development during unisexual reproduction is highly similar to that observed in canonical bisexual reproduction, except that the clamp cells along the hyphae are fused in bisexual reproduction but unfused in unisexual reproduction. Unisexual reproduction has also been observed in other heterothallic pathogenic fungal species, including *Candida albicans* and *Candida tropicalis* (33-36). While the molecular mechanisms underlying initiation and development of bisexual reproduction in *Cryptococcus* are well studied, how unisexual reproduction is initiated and regulated is less well understood. Nevertheless, studies have shown that unisexual reproduction involves a haploid to diploid transition that is likely achieved through endoreplication (37) and unisexual reproduction is promoted to a greater extent by the *MAT*α locus compared to *MAT***a** (38).

Recently, another dimorphic cellular transition, namely the formation of dramatically enlarged titan cells, has been identified and characterized in *Cryptococcus* species. While *Cryptococcus* typically grows as an encapsulated budding yeast (∼5 µm in diameter), titan cells are dramatically enlarged (>10 µm and up to more than 100 µm in diameter), polyploid (>2C and up to more than 128C in DNA content), and have cell wall and capsule characteristics distinct from those of standard yeast cells (39-44). The production of titan cells can be induced both *in vitro* and *in vivo*, and meiotic machinery has been implicated in regulating the ploidy transitions during this processes (45). Interestingly, titan cell formation (titanization) also involves sensing of a nutrient-poor environment, proper cell cycle control by cyclin-dependent kinases, and regulation by cAMP and MAPK pathways, similar to those regulating sexual reproduction (46-50). Biologically, both titan cell formation and sexual development involve changes in cellular polarization and cell cycle control. However, it is not known whether there are any connections between these two cellular morphological transitions.

In this study, we performed quantitative trait locus (QTL) analyses of a progeny population generated via sexual reproduction between two parental strains, one that is unisexually fertile and the other unisexually sterile. We identified two quantitative trait genes (QTGs), and the associated quantitative trait nucleotides (QTNs), that underlie the observed phenotypic variation in unisexuality. Both QTGs are involved in G-protein signaling, and we observed an epistatic interaction between the two QTGs and the *MAT*α locus that enhances unisexual hyphal growth. Interestingly, we identified an epistatic interaction between these QTGs, with one allele combination giving rise to a novel phenotype compared to either parent, in which large cells similar to titan cells are produced at high frequency under mating-inducing conditions. Further analyses showed that these large-cell forming progeny produced titan cells both *in vitro* and during infection in experimental animal models. Thus, our study provides insights into unisexual reproduction and titanization and evidence for an intersection between the regulation of these two morphological fates. Our findings also highlight the complexity of signaling pathways involved in controlling fungal cellular development, and the importance of inter-gene and inter-pathway interactions in mediating the phenotypic consequences of variation in such intricate regulatory networks.

## Results

### B3502 strains with the *RIC8* nonsense mutation undergo unisexual reproduction

There are four different stocks of B3502 in our lab collection (referred to here as strains B3502_A, B, C, and D), and they are copies of the stocks saved in the labs of Dr. Heitman (B3502_A; Duke University), Dr. Wiley (B3502_B), Dr. Vilgalys (B3502_C; Duke University), and Dr. Kwon-Chung (B3502_D; NIH/NIAID), respectively. Interestingly, when grown on MS solid medium (which induces mating and selfing) in the dark as a solo culture with no mating partner, both B3502_A and B3502_B produced extensive hyphae as well as basidia and basidiospore chains after prolonged incubation (∼4 weeks), while B3502_C and B3502_D produced smooth edges indicative of exclusive yeast-form growth (Figure 1A).

**Figure 1.**
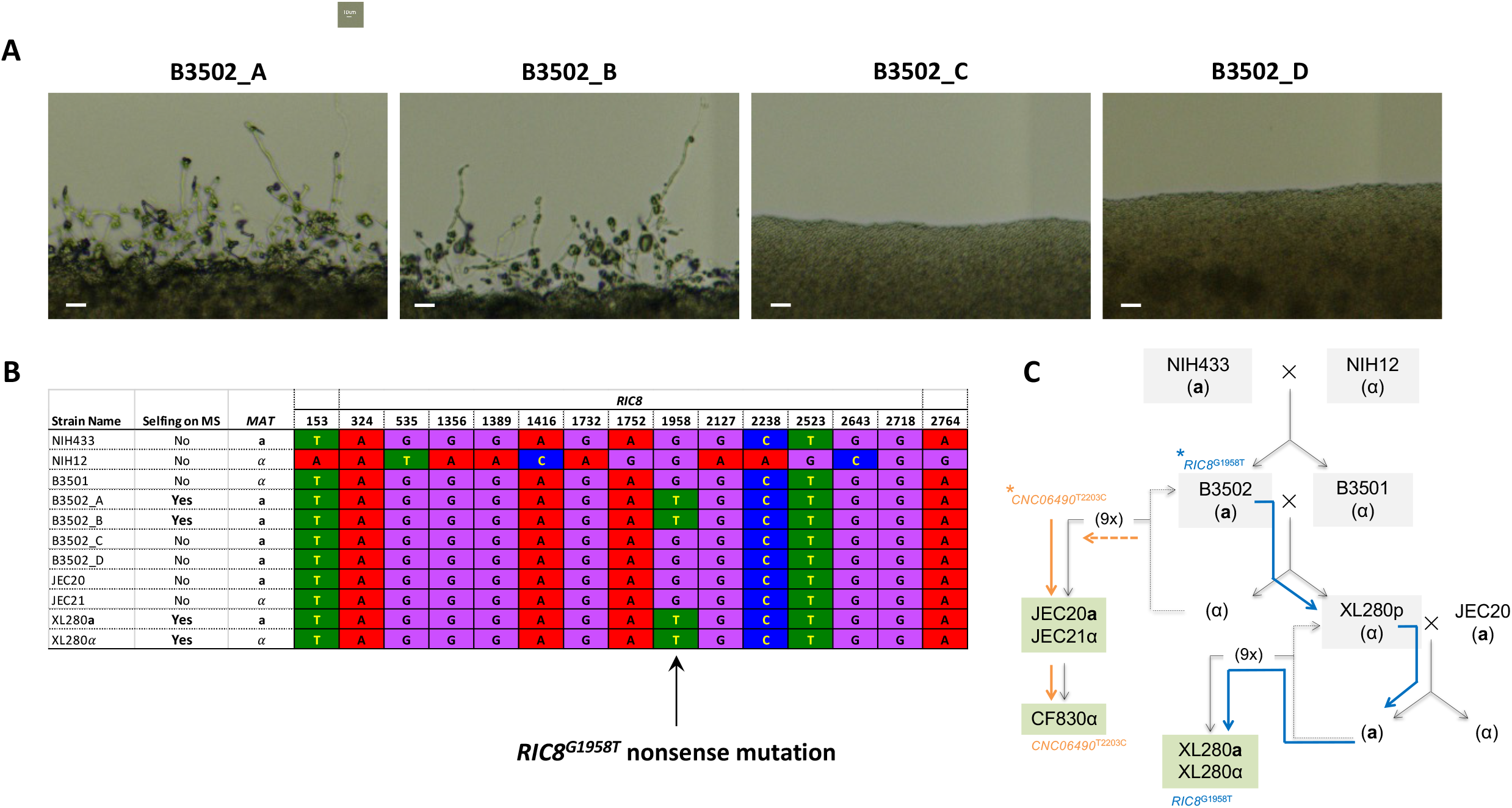
B3502 strains undergoing unisexual reproduction harbor a nonsense mutation in the *RIC8* gene. (A) Under mating-inducing conditions, only B3502_A and B3502_B undergo unisexual reproduction, while B3502_C and B3502_D do not. Scale bars represent 40 µm. (B) The *RIC8*^G1958T^ nonsense mutation is only present in strains capable of selfing, including B3502_A, B3502_B, as well as XL280**a** and XL280α that were derived from crosses involving B3502_B. (C) Genealogy of strains related to B3502. Blue and orange arrows trace the origin and inheritance of the QTGs influencing the phenotypes under mating condition among the B3502 stocks and the progeny from crosses B3502_A x CF830 and B3502_B x CF830.

B3502_B was one of the progenitor strains utilized in laboratory crosses to construct strain XL280, which undergoes robust unisexual reproduction (19). In a recent study, we identified a nonsense mutation in the *RIC8* gene (*RIC8*^G1958T^) of XL280 and found that this mutation underlies variation across several phenotypes linked to virulence, including tolerance to high temperatures, resistance to oxidative stress resistance, melanin production, and capsule size (51). Additionally, *RIC8* has been shown to interact with G-proteins involved in pheromone signaling during sexual reproduction in *C. neoformans* (52). Given these observations, we hypothesized that the robust selfing observed in the XL280 strain might be attributable to the *RIC8*^G1958T^ loss-of-function mutation.

To test this hypothesis, we analyzed the haplotypes generated from whole genome sequencing data of all four B3502 laboratory stocks, as well as for NIH12 and NIH433 that are the parents of the F_1_ sibling pair, B3501 and B3502 (Figure 1C)(53). By mapping the 114,223 genetic variants identified between the NIH12 and NIH433 genomes, we found that the four B3502 strains have almost identical haplotypes and are genetic recombinants of the NIH12 and NIH433 sequences, with very few segregating genetic variants. This is consistent with these strains being derived by lab passage from the same progenitor, which was a meiotic product of the cross between NIH12 and NIH433 (Supplemental Table S1; Supplemental Figures S1 and S2).

We next analyzed the selfing phenotype and the *RIC8* genotype of these strains, together with a collection of genealogically closely-related strains, including JEC20, JEC21, XL280**a**, and XL280α (Figure 1B). We found that the *RIC8*^G1958T^ allele is not present in either of the two parental strains (NIH12 and NIH433) of B3502. Additionally, we found that while the *RIC8*^G1958T^ allele is present in all strains capable of selfing, including B3502_A, B3502_B, and the XL280**a** and XL280α congenic strains that are genetic descendants of B3502_B, the *ric8* nonsense mutation was absent in all other strains analyzed here. This is consistent with the previous observation that the nonsense mutant *RIC8*^G1958T^ allele originated from a *de novo* mutation within a select few B3502 strains, and was subsequently inherited by the XL280 congenic strains (Figure 1C)(51). The observation that the *RIC8*^G1958T^ allele is only present in the strains that undergo robust selfing indicates this allele might be associated with unisexual reproduction in *Cryptococcus deneoformans*.

### Progeny from B3502 crosses produced segregating phenotypes under mating conditions

To further investigate whether the *RIC8*^G1958T^ nonsense mutation is associated with the selfing phenotype observed in B3502_A and B3502_B, we conducted crosses between each of the four B3502 strains and strain CF830 (JEC21, *MAT*α *NOP1-GFP-NAT*), recovered meiotic progeny by dissecting the spores produced from these crosses, assayed the selfing phenotypes of the progeny on solid MS medium, and genotyped the progeny for their mating types (*MAT***a** vs. *MAT*α) and the *RIC8* allele type (WT vs. *RIC8*^G1958T^ nonsense mutant) (Supplemental Tables S2-S3; Supplemental Figure S3). Because strain CF830 does not undergo selfing and grows as yeast under mating conditions, if the *RIC8*^G1958T^ allele is responsible for the selfing phenotype, we should expect: 1) only crosses involving B3502_A and B3502_B to generate progeny capable of selfing; 2) among progeny recovered from crosses involving B3502_A and B3502_B, the ratio between selfing and non-selfing phenotypes should be approximately 1:1; and 3) the progeny that are able to undergo selfing should all inherit the *RIC8*^G1958T^ allele from the B3502_A and B3502_B parents.

Consistent with this hypothesis, the vast majority of the progeny dissected from crosses B3502_C x CF830 (47 of 47) and B3502_D x CF830 (21 of 24) were not able to undergo selfing and grew as yeasts under mating conditions (Table 1). The only exceptions were three progeny dissected from the crosses B3502_D x CF830 that showed a transgressive selfing phenotype. Subsequent analyses indicate these progeny inherited both **a** and α *MAT* alleles, and thus were likely diploid or aneuploid and capable of undergoing bisexual reproduction under mating-inducing conditions (Supplemental Tables S4-S5).

**Table 1.**
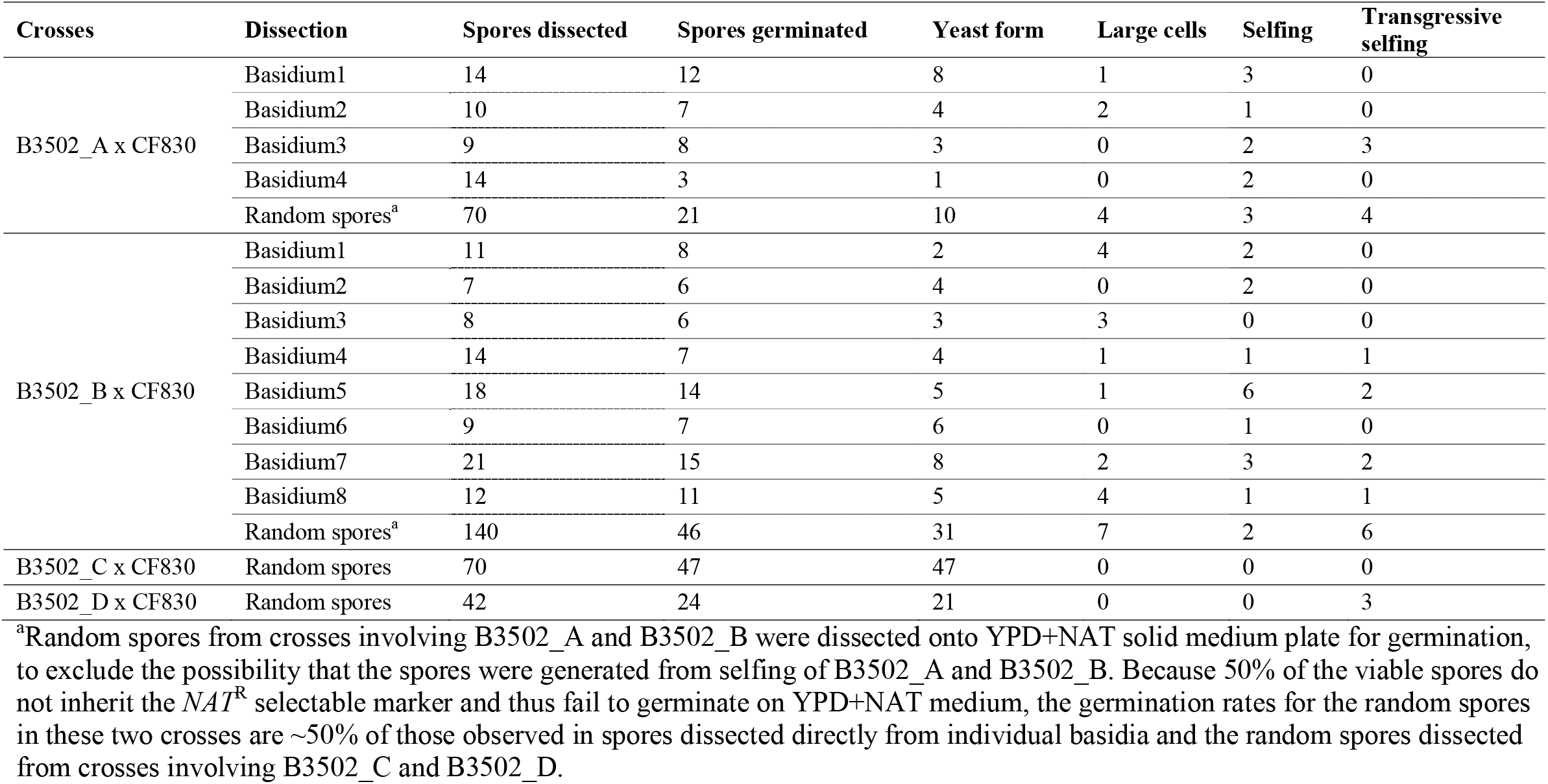
Summary of the phenotypes under mating condition among progeny produced by crosses involving B3502 strains

| Crosses | Dissection | Spores dissected | Spores germinated | Yeast form | Large cells | Selfing | Transgressive selfing |
| --- | --- | --- | --- | --- | --- | --- | --- |
| B3502_A x CF830 | Basidium1 | 14 | 12 | 8 | 1 | 3 | 0 |
|  | Basidium2 | 10 | 7 | 4 | 2 | 1 | 0 |
|  | Basidium3 | 9 | 8 | 3 | 0 | 2 | 3 |
|  | Basidium4 | 14 | 3 | 1 | 0 | 2 | 0 |
|  | Random spores<sup>a</sup> | 70 | 21 | 10 | 4 | 3 | 4 |
| B3502_B x CF830 | Basidium1 | 11 | 8 | 2 | 4 | 2 | 0 |
|  | Basidium2 | 7 | 6 | 4 | 0 | 2 | 0 |
|  | Basidium3 | 8 | 6 | 3 | 3 | 0 | 0 |
|  | Basidium4 | 14 | 7 | 4 | 1 | 1 | 1 |
|  | Basidium5 | 18 | 14 | 5 | 1 | 6 | 2 |
|  | Basidium6 | 9 | 7 | 6 | 0 | 1 | 0 |
|  | Basidium7 | 21 | 15 | 8 | 2 | 3 | 2 |
|  | Basidium8 | 12 | 11 | 5 | 4 | 1 | 1 |
|  | Random spores<sup>a</sup> | 140 | 46 | 31 | 7 | 2 | 6 |
| B3502_C x CF830 | Random spores | 70 | 47 | 47 | 0 | 0 | 0 |
| B3502_D x CF830 | Random spores | 42 | 24 | 21 | 0 | 0 | 3 |

<sup>a</sup>Random spores from crosses involving B3502\_A and B3502\_B were dissected onto YPD+NAT solid medium plate for germination, to exclude the possibility that the spores were generated from selfing of B3502\_A and B3502\_B. Because 50% of the viable spores do not inherit the *NAT<sup>R</sup>* selectable marker and thus fail to germinate on YPD+NAT medium, the germination rates for the random spores in these two crosses are ~50% of those observed in spores dissected directly from individual basidia and the random spores dissected from crosses involving B3502\_C and B3502\_D.

From crosses between B3502_A and CF830, we dissected spores from four individual basidia, with spore germination rates ranging from 21% (basidium 4) to 89% (basidium 3) (Table 1). We also dissected 70 random spores, of which 21 (30%) germinated on YPD+NAT solid medium, indicating theoretically the germination rates among the random spores should be ∼60%, which is within the range of germination rates observed among individual basidia. Similarly, we dissected eight individual basidia and 140 random basidiospores from crosses between B3502_B and CF830, with germination rates ranging from 50% to 92% (average ∼74%).

To our surprise, when spores from crosses B3502_A x CF830 and B3502_B x CF830 were recovered and incubated under selfing-inducing conditions, the progeny exhibited four different phenotypes: 1) yeast form, similar to the CF830 parent with a smooth colony edge and surface, and mostly yeast cells in the population when inspected by microscopy; 2) large cells, which manifest rough surfaces and uneven edges of the colony, and whose cell population is composed of normal yeast cells mixed with large cells up to >30 µm in diameter; 3) selfing, as seen in the B3502_A and B3502_B parental strains with robust hyphae produced at the edge of the colony, and with both yeast cells and hyphae when observed by microscopy; and 4) transgressive selfing, in which longer and more dense hyphae were produced at the colony periphery, and considerably more and longer hyphae observed by microscopy (Table 1 and Figure 2). These findings suggest that more than one genetic locus contributes to the selfing phenotype of strains B3502_A and B3502_B.

**Figure 2.**
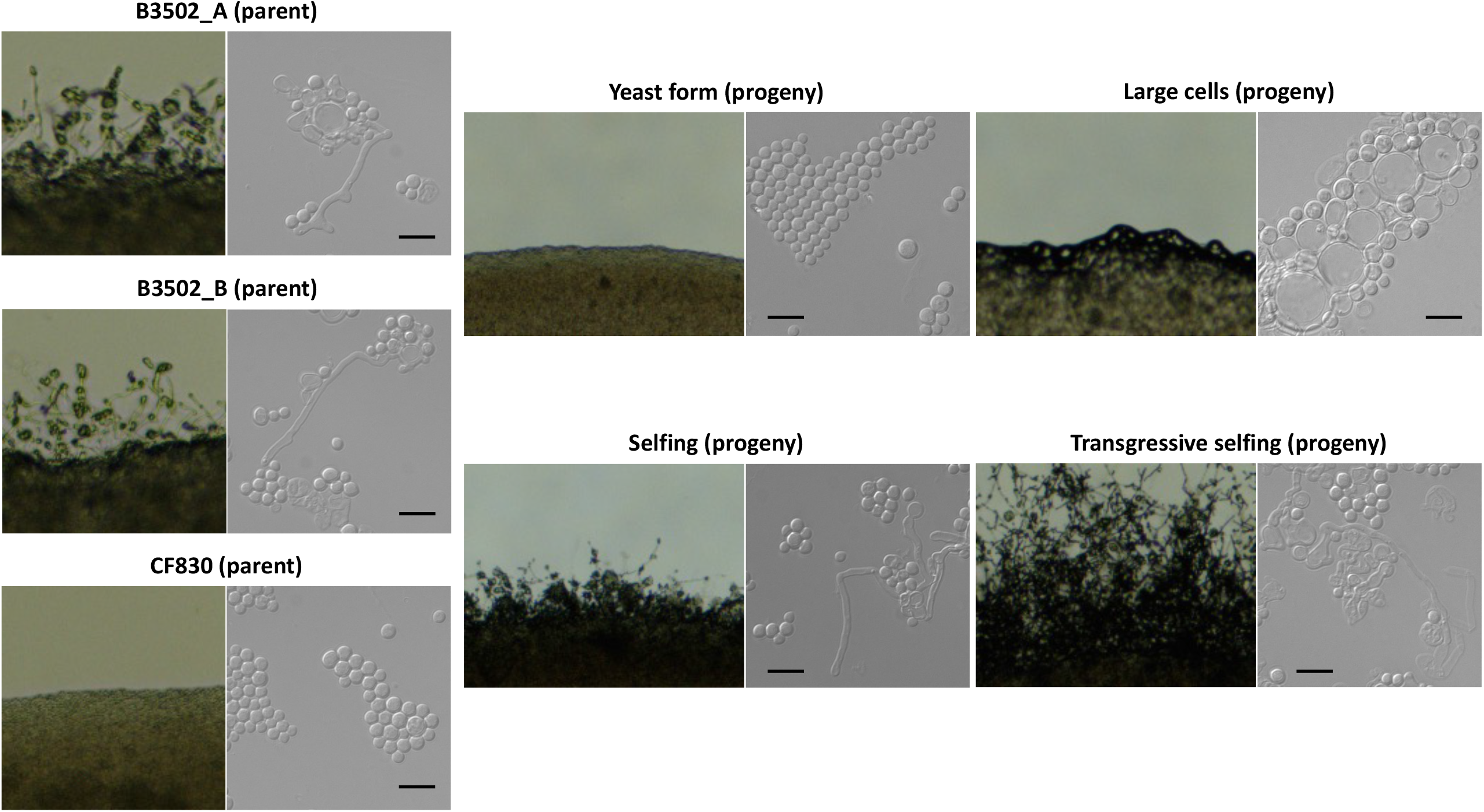
Colony and cellular morphologies under mating conditions of progeny recovered from B3502 crosses. Images of solo cultures on MS solid medium under mating condition (left) and cells under microscope (right) are shown for the three parental strains (B3502_A, B3502_B, and CF830) and four progeny representing four different phenotypes under mating conditions. Scale bar represents 10 µm.

We genotyped progeny from the B3502_A and B3502_B crosses for both the *MAT* locus and the *RIC8* gene. Our goal was to examine the relationship between the genotypes at these loci and the observed cellular phenotypes in this progeny population. For the B3502_A crosses, of the 24 progeny that grew as yeasts under selfing-inducing conditions, all possessed a wild-type *RIC8* allele. In contrast, of the 24 progeny that did not grow as yeasts under selfing-inducing conditions (large cells or selfing), all but one had the *RIC8*^G1958T^ nonsense allele (Table 2). This observation was further supported by the analysis of spores recovered from the B3502_B crosses. Specifically, nearly all of the progeny (52 of 57) that grew as yeast had the wildtype *RIC8* allele, and all the progeny (60 of 60) that did not grow as yeasts had the *RIC8*^G1958T^ nonsense mutant allele (Table 2).

**Table 2.** Summary of association between phenotypes under mating conditions and genotypes at the *RIC8* locus

|  | Crosses | No. of spores analyzed | Yeast form |  | Non-yeast form<br>(large-cell, selfing, transgressive selfing) |  |
| --- | --- | --- | --- | --- | --- | --- |
|  |  |  | *RIC8*_WT | *RIC8*_nonsense | *RIC8*_WT | *RIC8*_nonsense |
| Basidium-specific<br>dissection | B3502A x CF830 | 30 | 16 | 0 | 0 | 14 |
|  | B3502B x CF830 | 71 | 36 | 0 | 0 | 35 |
| Random spore<br>dissection | B3502A x CF830 | 18 | 8 | 0 | 1 | 9 |
|  | B3502B x CF830 | 46 | 16 | 5 | 0 | 25 |
|  | B3502C x CF830 | 47 | 47 | 0 | 0 | 0 |
|  | B3502D x CF830 | 24 | 21 | 0 | 3 | 0 |

Thus, our results suggest there is a near 100% co-segregation between the *RIC8* wild-type allele and yeast-form growth under selfing-inducing conditions, which is consistent with our hypothesis. However, progeny with the *RIC8*^G1958T^ nonsense mutant allele do not always exhibit the selfing phenotype as observed in the B3502 parent. Instead, their phenotypes vary among large cells, selfing, and transgressive selfing, suggesting there are additional genetic factors in the genome that act epistatically with the *ric8* nonsense mutation to determine the non-yeast form phenotype in these progeny.

Interestingly, of the 19 progeny that displayed a transgressive selfing phenotype, all had the α allele at the *MAT* locus (Table 1; Supplemental Tables S2 and S3), which is significantly different from the expected frequency of the mating types if it were to be distributed randomly (*p-value* < 0.05, Fisher’s Exact Test). Thus, the *MAT*α allele is necessary but not sufficient for the transgressive selfing phenotype in the presence of the *RIC8*^G1958T^ nonsense mutation (also see QTL analyses below), which again suggests the presence of additional, yet to be identified loci in the genome that are contributing to the large cell, selfing, and transgressive selfing phenotypes.

### Large-cell progeny produced titan cells *in vitro* and *in vivo*

We next investigated whether the large cells produced by the large-cell progeny have characteristics of titan cells. When grown under titan-cell inducing conditions (46), of the four phenotypes identified under mating-inducing conditions, only the large-cell progeny produced enlarged cells at significant frequencies when inspected by microscopy (Figure 3A). Analysis of the cell size distributions showed that the large-cell progeny produced cells with diameters that are larger than 10 µm at frequencies considerably higher than the two parental strains and similar to the H99 *usv101*Δ mutant strain, which is known to produce titan cells at an elevated frequency (Figure 3B and Supplemental Figure S4). Additionally, FACS analyses of the cells grown under *in vitro* titan-cell inducing conditions also revealed an increase in polyploid cells in the large-cell progeny, but not in the two parental strains (Figure 3C). Finally, while the two parental strains remained as yeast cells in the lungs of infected mice, the large-cell progeny produced enlarged cells with thickened capsules in the lungs at 8 days post-inoculation (Figure 3D). Taken together, our analyses provided substantial evidence that these large-cell progeny produced *bona fide* titan cells *in vitro* and in *vivo*.

**Figure 3.**
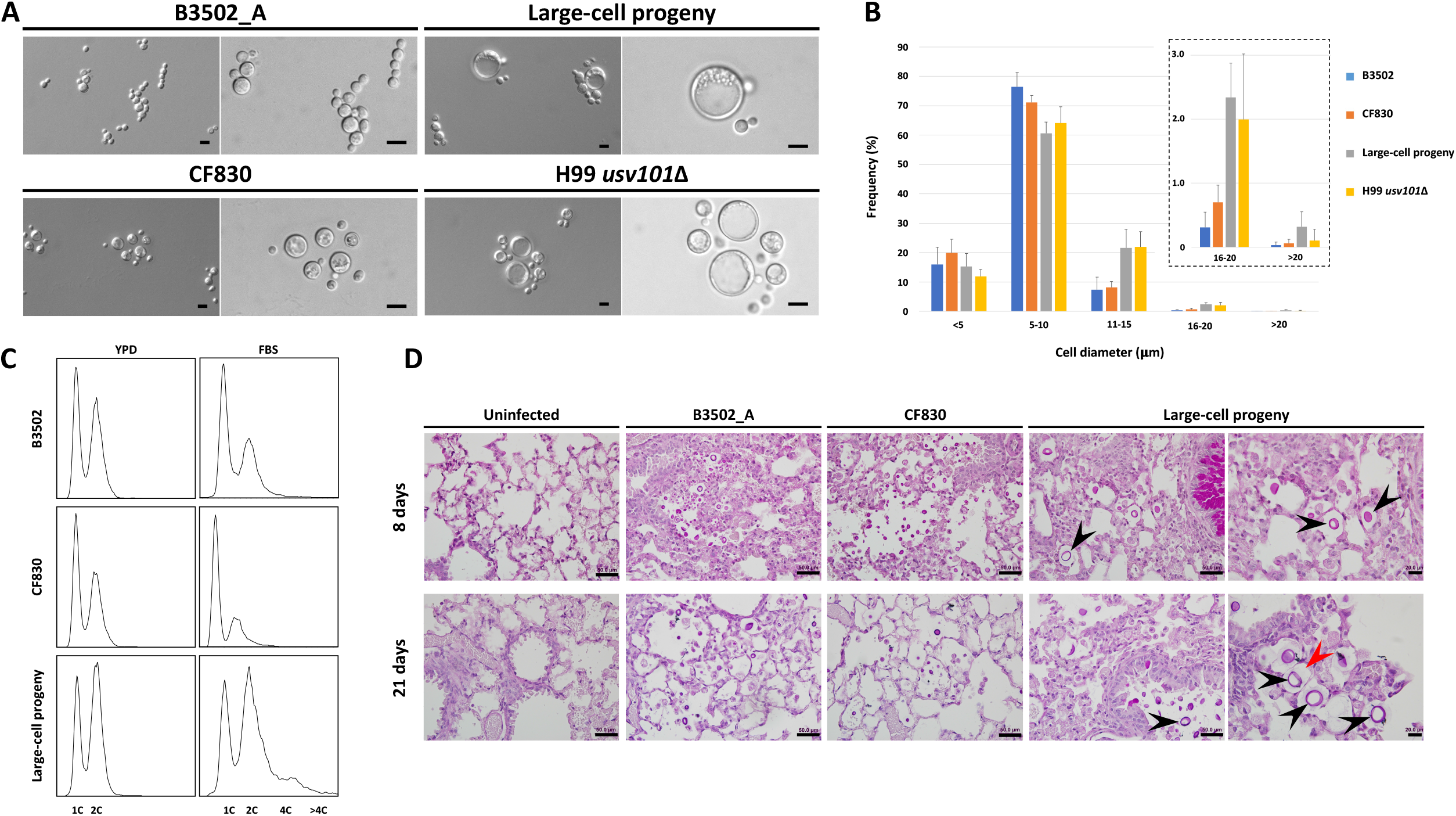
Large-cell progeny produce titan cells *in vitro* and *in vivo*. A) Large cells are produced by the large-cell progeny under *in vitro* titan cell-inducing conditions. For each strain, the images on the left and right are taken using 20x and 40x objectives, respectively. The scale bars represent 10 µm. Strain H99 *usv101*Δ is known to produce titan cells and is included as positive control. (B) Under *in vitro* titan cell inducing conditions, the large-cell progeny produced cells with diameters > 10 µm at higher frequencies. The inset shows frequencies of cells with diameters >15 µm using an adjusted Y-axis. More than 1000 cells from each strain were analyzed using Cellometer and independent measurements of cell sizes of each strain using ImageJ are presented in Supplemental Figure S4. (C) FACS analysis showed that large-cell progeny produced cells with elevated ploidy under *in vitro* titan cell-inducing conditions. The x- and y-axes show DNA content levels and cell counts, respectively. (D) Large-cell progeny produced enlarged cells *in vivo*. The top and bottom panels are histopathology images of mouse lungs dissected 8 and 21 days after inoculation, respectively. For large-cell progeny, the images on the left are at the same scale as those of the other strains (scale bar = 50 µm), while the images on the right are taken at higher magnification (scale bar = 20 µm). The black arrowheads highlight enlarged cells observed in the lungs inoculated with large-cell progeny; the red arrowhead highlights the thickened capsule surrounding the enlarged cell.

### QTL analyses identified an additional candidate gene underlying the large-cell phenotype

To further investigate other genetic factors contributing to the diverse phenotypes observed among the F_1_ progeny, we utilized Illumina whole genome sequencing from 70 spores dissected from crosses B3502_A x CF830 and B3502_B x CF830 (Supplemental Tables S2-S3) and identified 1,814 high quality genetic variants segregating among these progeny. Of these 1,814 bi-allelic variants, 60 (∼3%) are randomly distributed across the genome, while the remaining 1,754 variants group into three distinct clusters: one contains 1,286 variants (71%) and corresponds to the *MAT* locus (i.e. genetic variation between the *MAT***a** and *MAT*α alleles), and the other two are located on the left- and right-end of Chromosome 2 and contain 252 (∼14%) and 216 (∼12%) variants, respectively (Supplemental Figure S1). We then conducted QTL mapping using these variants together with the cellular phenotypes under mating-inducing conditions of the 70 progeny and the three parental strains.

For the initial QTL analysis, we classified the progeny phenotypes into binary scores as yeast form and non-yeast form, with the latter including large-cell, selfing, and transgressive selfing phenotypes (Supplemental Figure S5A). With this binary characterization of phenotypes and using an information-theoretic approach based on mutual information (54), we identified two QTL loci on Chromosome 14: a QTL of modest effect (I (*G*_*n*_; *S*) = 0.184 nats) at approximately 134 kb and a QTL with large effect (I (*G*_*n*_; *S*) = 0.581 nats) at approximately 388.8 kb (Figure 4A, Supplemental Figure S5B). The first QTL maps to a SNP (G ⟶ C) within the first exon of the gene *CNN00400*, which is a conserved hypothetical protein in the *C. deneoformans* reference genome. This gene has an annotated ortholog, *IRK7*, in the reference genome of the sibling species *C. neoformans* that encodes a Ca2+/ calmodulin-dependent protein kinase. The *CNN00400*^G1740C^ mutation is predicted to cause a nonsynonymous change – glycine to arginine – in the protein product (Supplemental Figure S6). The second, major effect QTL corresponds to the previously described *RIC8*^G1958T^ nonsense mutation variant within the penultimate exon at 1,958 bp within this gene (Figure 4A).

**Figure 4.**
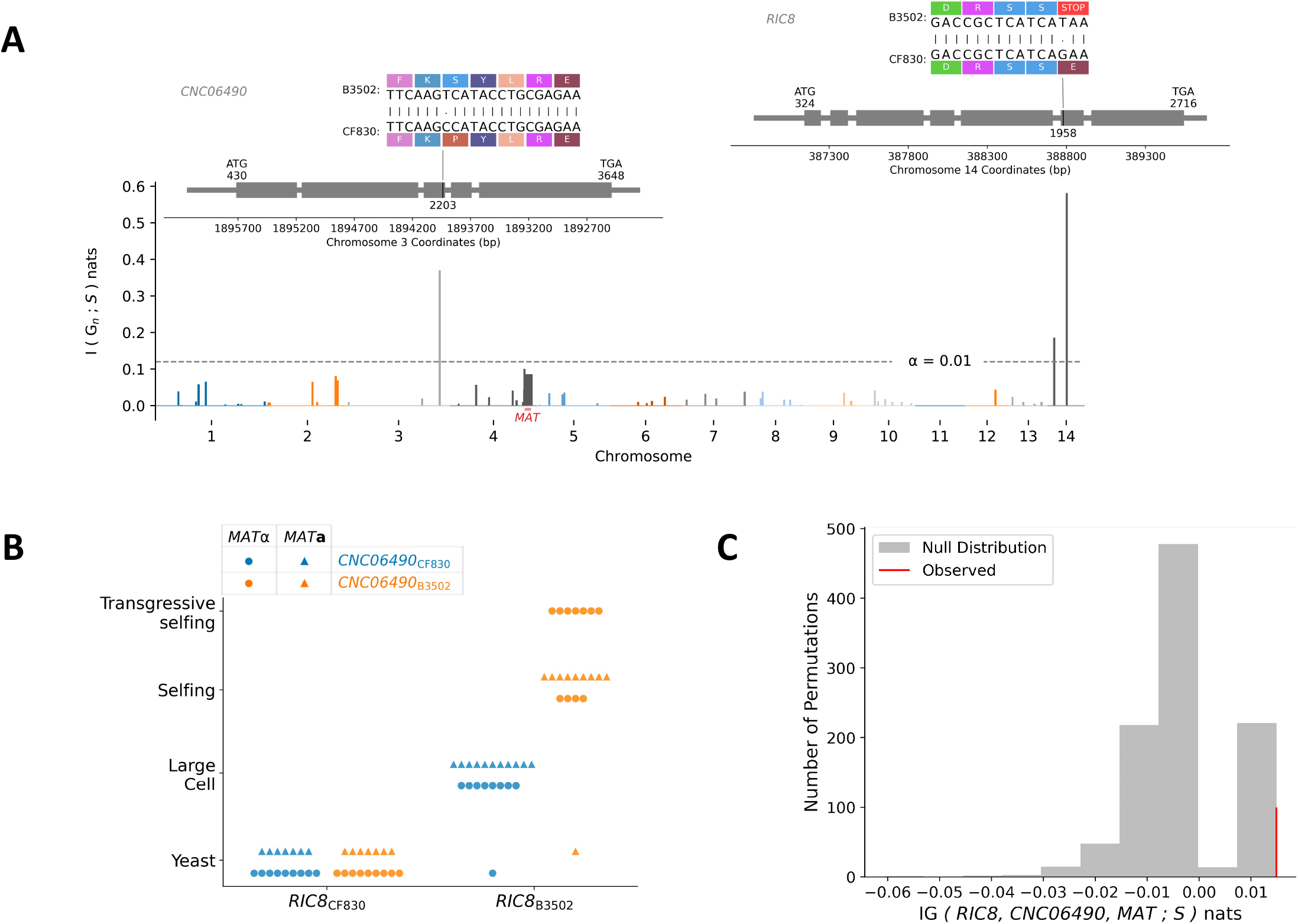
QTL analyses identify two QTGs associated with selfing and large cell phenotypes. (A) Manhattan plot displaying the genome-wide strength in association (measured with a mutual information statistic (I (*Gn* ; *S*) - y-axis) between genetic variants (x-axis) and the selfing-cellular phenotypes. Colors separate chromosomes, and the position of the *MAT* locus is indicated by a red horizontal bar and text. Above the QTL on Chromosomes 3 and 14 are gene models for the QTGs, *CNC06490* and *RIC8*, displaying the position and predicted effects of the identified QTN within each gene (left and right respectively). (B) A phenotype by genotype plot depicting the cellular phenotypes (y-axis) of the F_1_ progeny as a function of the parental *RIC8* allele (x-axis) colored by the allele inherited at *CNC06490*. Dots and triangles denote the presence of the *MAT*α *or MAT***a** allele for each progeny, respectively. C) Information gain null distribution (grey) and observed informational gain statistic (red) testing for a three-way, epistatic interaction between the QTN (*RIC8*^G1958T^ and *CNC06490*^T2203C^) and the *MAT* locus.

Further analyses showed that the two QTL on Chromosome 14 are highly linked (Spearman ρ = 0.583, *p*-value < 0.01) with a recombination event occurring between these loci in only 21% (15/70) of the progeny. Treating these two QTL peaks along Chromosome 14 as the same QTL and constructing a 99% confidence interval via bootstrapping centered the effect on the *RIC8*^G1958T^ mutation. Given the dramatic, predicted change in protein sequence of the *RIC8*^G1958T^ allele, together with our results from earlier analyses based on the *RIC8* genotyping, that is, the nonsense mutant allele *RIC8*^G1958T^ is necessary for the non-yeast form growth under mating conditions, we therefore labeled the *RIC8*^G1958T^ mutation as the QTN underlying this binary phenotype.

We next divided the non-yeast form phenotype into two groups: large cells and selfing (including both selfing and transgressive selfing) (Supplemental Figure S5A) and repeated the QTL mapping with this “ternary” phenotypic classification. In this analysis, we detected the same QTL identified in the binary analysis – which included the QTN, *RIC8*^G1958T^, on Chromosome 14 – and a second, novel large-effect QTL (I (*G*_*n*_; *S*) = 0.37 nats) on Chromosome 3 (Figure 4A, Supplemental Figure S5B). This QTL on Chromosome 3 maps to a nonsynonymous substitution in the third exon of the gene *CNC06490*, changing a proline to a serine in strains B3502_A and B3502_B (Figure 4A). In the *C. deneoformans* reference genome, the gene *CNC06490* is annotated as a putative Rho GTPase-activating protein (RhoGAP).

Among the strains that are closely related to B3502, the *CNC06490*^2203C^ allele is only present in JEC20, JEC21, and CF830 that is derived from JEC21, while all other strains possess the *CNC06490*^2203T^ allele (Figure 1C), suggesting that the serine codon at the 2203 bp location is the ancestral state, and the T ⟶ C substitution arose prior to the backcrossing conducted to generate the JEC20 and JEC21 congenic strains.

The fact that the *CNC0649*0 QTL was only detectable when the large-cell phenotype was treated as a separate phenotypic class suggests that this QTN underlies the cellular morphology of cells growing as large cells under mating conditions. Additionally, our results suggest that the two QTN, corresponding to the *CNC06490*^T2203C^ and *RIC8*^G1958T^ mutations, respectively, interact and collectively contribute to the large-cell phenotype (Figure 4B). Indeed, when the genotypes of the progeny at these two loci were analyzed together with their phenotype, we found again that all progeny that inherited the *RIC8* wild-type allele from the CF830 parent had the yeast-form phenotype. However, when the progeny inherited the *RIC8*^G1958T^ nonsense mutant allele from the B3502 parent, they exhibited a selfing (selfing and transgressive selfing) phenotype when they also inherited the *CNC0649*0 B3502 allele, but a large cell phenotype when they inherited the *CNC06490* CF830 allele. Taken together, our analyses suggest an epistatic interaction between the *CNC06490* and *RIC8* genes that determines the morphological fates of *C. deneoformans* cells grown under mating-inducing conditions.

Finally, we repeated the QTL analyses with four different phenotypic classes: yeast form, large cell, selfing, and transgressive selfing (Supplemental Figure S5A). In this scenario, we identified the same QTN (*CNC06490*^T2203C^ and *RIC8*^G1958T^) as in the ternary analysis (Figure 4A, Supplemental Figure S5B). Moreover, compared to the ternary analysis, we observed an enhanced signal corresponding to the *MAT* locus when transgressive selfing was analyzed as a separate phenotypic group, further suggesting the *MAT*α allele contributes to the transgressive selfing phenotype (Figure 4B, Supplementary Figure S4B).

To determine whether there is strong evidence for three-way interactions between *RIC8, CNC06490*, and the *MAT* locus, we carried out an analysis of epistasis based on a measure of “informational gain” among these loci using the four cellular phenotypic states (51). This analysis identified a significant interaction between the two QTN (*RIC8*^G1958T^ and *CNC06490*^T2203C^) and the *MAT* locus (IG (*RIC8, CNC06490, MAT*; S) = 0.015 nats, *p*-value < 10^−10^; Figure 4C). This was consistent with our previous observations based on the progeny genotypes where all progeny exhibiting the transgressive selfing phenotype also inherited the α mating type (Supplemental Tables S2 – S5), indicating the *MAT*α allele is necessary but not sufficient for the transgressive selfing phenotype in the presence of the *RIC8*^G1958T^ nonsense mutation. This result is also consistent with previous studies identifying the *MAT*α allele as a prominent QTL for hyphal growth during selfing (38).

### Deletion of the *RIC8* gene leads to attenuated mating in bilateral crosses in *C. neoformans*

We identified a strong correlation between the *RIC8*^G1958T^ nonsense mutation and the non-yeast form growth phenotypes under mating-inducing conditions in *C. deneoformans*. We next tested whether this genetic change produces the same phenotype in the sister species, *C. neoformans*. To accomplish this, we deleted the *RIC8* gene in H99 and KN99**a**, two congenic *C. neoformans* strains of opposite mating types. Under mating-inducing conditions these *ric8*Δ strains grew as yeasts, which is different from the phenotypes observed in *C. deneoformans* B3502_A and B3502_B strains. This suggests that the regulation of the cellular phenotype by the *RIC8* gene may differ between these two sibling species, or that there are strain- or background-specific effects.

Interestingly, when we tested these *ric8*Δ strains in bisexual mating assays, we found that while unilateral crosses did not show any discernible differences compared to the wild-type crosses, the bilateral crosses in which the *RIC8* gene is absent in both parents showed attenuated hyphal growth and highly reduced production of basidia and basidiospores, suggesting a functional *RIC8* gene is required for robust bisexual reproduction in *C. neoformans* (Figure 5).

**Figure 5.**
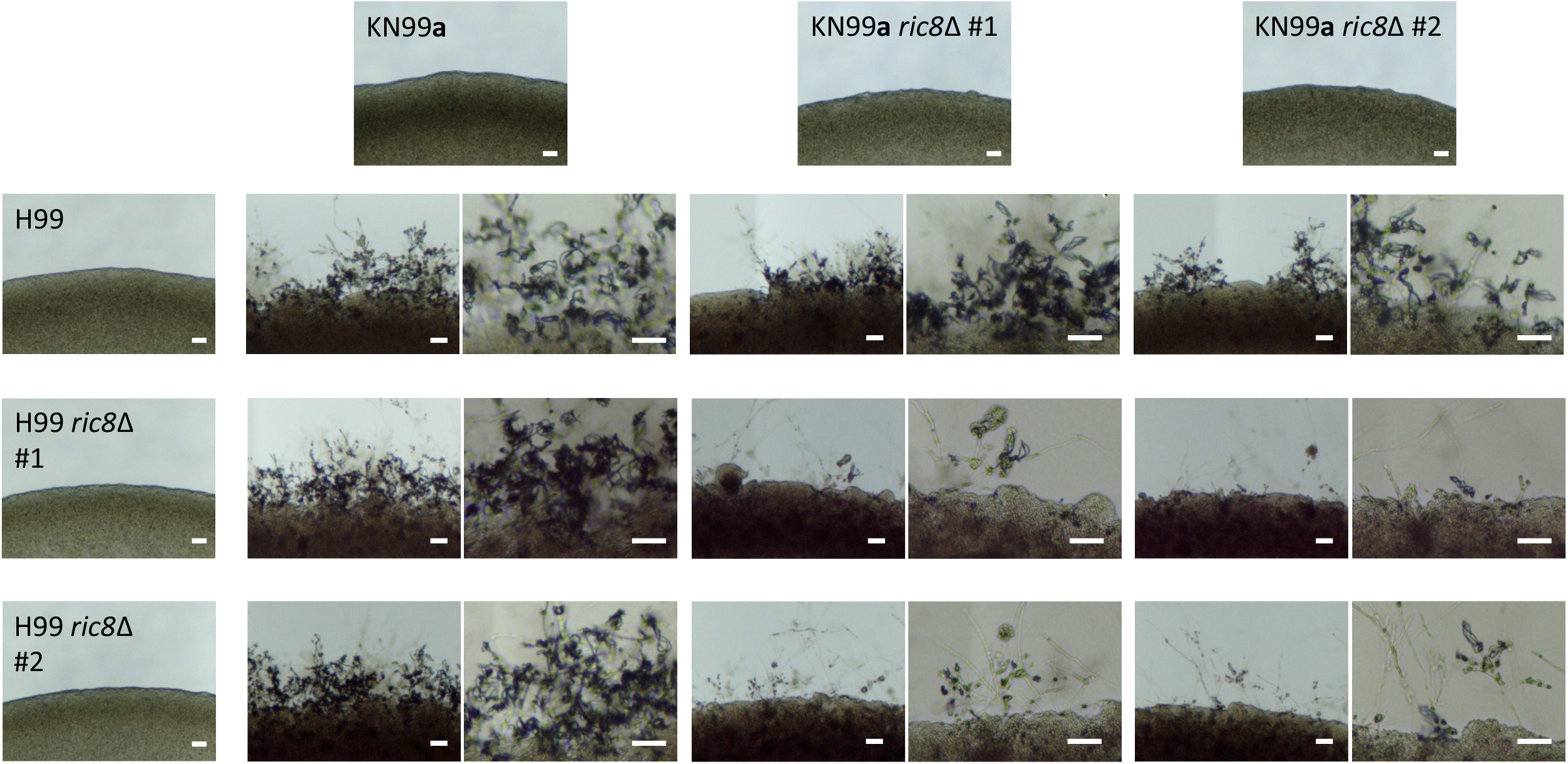
Deletion of *RIC8* leads to attenuated bi-sexual mating in *C. neoformans*. Images of mating co-cultures at lower (left) and higher (right) magnifications are shown for the wild-type control cross between H99 and KN99**a**, as well as reciprocal unilateral crosses and bilateral crosses of two *ric8*Δ deletion strains each in the H99 and KN99**a** congenic backgrounds. The images in the top row and on the far left are solo cultures of the parental strains at lower magnifications. While all four unilateral crosses showed mating efficiencies that are comparable to the wild-type control cross, the four *ric8*Δ deletion bilateral crosses showed dramatically less robust mating, manifested as reduced hyphal growth and fewer basidia and basidiospores. Scale bars represent 40 µm.

## Discussion

We conducted QTL analysis of the selfing phenotype observed in *C. deneoformans* strains in progeny generated from crosses between selfing and non-selfing strains, and we identified two major QTGs that govern the variation of several cellular morphologies under mating-inducing conditions. Interestingly, both QTGs appear to be related to the regulation of G proteins, and the epistatic interaction between these two loci underlies a novel large-cell phenotype observed in a subset of the progeny analyzed here.

G proteins and their signaling pathways are integral in a variety of cellular processes in eukaryotes including fungi, such as sensing stimuli in the surrounding environment, transduction of these signals inside the cell, and regulating cellular activities and development (55). In *Cryptococcus*, G proteins play important roles in cell cycle control, sensing and adapting to environmental stresses, mating and sexual development, as well as virulence and pathogenicity (56). The *RIC8* QTG identified in this study encodes a regulator of the cAMP-PKA signaling pathway that transmits G-protein signals inside the cell and has been shown in a recent QTL study to be highly pleiotropic, contributing to variation in four virulence related traits: melanization, capsule size, thermal tolerance, and resistance to oxidative stress in *C. deneoformans* (51). In *C. neoformans* Ric8 is known to interact with the G-protein α subunits Gpa1 and Gpa2 involved in pheromone responses and mating differentiation (52). In our study, we found that a fully functional *RIC8* allele is a critical, positive regulator for *C. deneoformans* growth as a yeast under conditions that typically induce selfing and sexual reproduction. It is possible that in the absence of *RIC8*, the transduction of G-protein signals, such as those involved in pheromone production and pheromone response, becomes compromised, triggering cells to initiate unisexual development in the absence of compatible mating partners for bisexual reproduction.

The second major QTG identified in this study, *CNC06490*, encodes a putative Rho GTPase-activating protein (GAP), and it only affects the phenotypic outcomes of progeny in the presence of the *RIC8*^G1958T^ nonsense mutant allele, indicative of an epistatic interaction between alleles of the two genes. This result echoes findings from a recent QTL study of virulence and stress response-related phenotypes in *C. deneoformans*, which found a variety of phenotypic outcomes result from epistatic interactions between the QTLs identified (51). Rho GTPases coordinate a wide variety of cellular processes in eukaryotes, including cell polarity and cell cycle progression (57). Rho GTPases such as Ras, Cdc42 paralogs, and Rac paralogs are involved in polarized growth, cytokinesis, cell cycle progression, stress response, and virulence of pathogenic fungi including *Candida albicans* and *C. neoformans* (58-64).

In the presence of the *RIC8*^G1958T^ nonsense mutation, having the *CNC06490* allele from the CF830 parent allowed progeny to form large cells at high frequencies under mating-inducing conditions. These large cells resemble *C. neoformans* titan cells in their size and also appear to have thickened cell walls that hinder staining of nuclei in live cells with Hoechst nuclear dye.

Additionally, we found that: i) *C. deneoformans* forms large cells under mating-inducing conditions, which are nutrient poor; ii) this large-cell phenotype involves a major QTG corresponding to a gene encoding GAP regulator of Rho GTPase; and iii) the large-cell phenotype requires epistatic interactions between the GTPase GAP major QTG and *RIC8*, which encodes a regulator of the cAMP-PKA signaling pathway. These findings are consistent with recent studies showing that titanization in *Cryptococcus* involves exposure to nutrient-limited environments, regulation by G proteins, and cAMP signaling (39-41, 46-48). Our further characterization of the large-cell forming progeny showed that they indeed form titan cells both *in vitro* under titan-cell inducing conditions and *in vivo* in the context of the infected murine lung. Thus, these large-cell progeny could be utilized to screen for positive regulators of titanization in *Cryptococcus* through approaches such as random insertional mutagenesis.

It is interesting that the two major QTGs identified in this study, *RIC8* and *CNC06490*, are predicted to have opposite functions on G proteins. While the *RIC8* gene, encoding a guanine nucleotide exchange factor (GEF), functions to enhance G protein signaling, the *CNC06490* gene encodes a GTPase-activating protein (GAP) that limits the activation of G proteins by terminating the G-protein signaling event. It is not clear which G proteins these two QTGs directly interact with to regulate the phenotypic variation observed in this study, although it is known that Ric8 interacts with Gpa1 and Gpa2 (52). Given the detected epistatic interaction between *RIC8* and *CNC06490*, it is possible there is overlap in the G proteins or G-protein signal transduction pathways impacted by these two QTGs.

Sexual development and the associated changes in cellular morphology are complex processes involving multiple genes and genetic interactions. We identified two major QTGs (and their associated QTNs) that interact epistatically with each other and the *MAT* locus to regulate variation in cellular morphologies in *C. deneoformans* under mating-inducing conditions. Such complexities in cellular morphology and their regulation could be advantageous for *Cryptococcus* to survive in the environment where they are faced with a variety of pressures, including nutrient limitation and predation by amoeba, as well as from attacks by the immune system and exposure to antifungal drugs when infecting human hosts (65). It is clear from our data that additional genetic loci influence these phenotypes that have yet to be identified. For example, five progeny that inherited the *RIC8*^G1958T^ nonsense mutant allele yet still grew as yeasts on MS medium (Table 2). This result could be due to the existence of modifiers of the *ric8* nonsense allele segregating in these progeny. Additional studies are needed to define these unknown contributing and interacting genetic factors, possibly by analyzing additional meiotic progeny from these crosses or conducting backcrosses to increase the representation of interacting allelic combinations and gain sufficient statistical power.

Interestingly, neither of the two QTGs identified here corresponds to the five QTLs identified in a previous study of variation in selfing and hyphal development in *C. deneoformans* (38), further highlighting the complexity of the genetic factors involved in *Cryptococcus* sexual development. The two parental strains used to generate the mapping population in that previous study were both self-filamentous, and our analysis confirmed that both contained the *ric8* nonsense mutation, explaining why the *RIC8* gene was not detected as one of the major QTLs.

Nevertheless, the *MAT* locus was identified as one of the major QTLs and the α allele was found to enhance filamentation (38), which is consistent with our finding that all of the progeny that showed a transgressive selfing phenotype harbor the *MAT*α allele. The **a** and α *MAT* alleles differ significantly both at the nucleotide level as well as structurally, with extensive chromosomal rearrangements including translocations and inversions (31, 32, 66). A variety of phenotypes have been shown to be influenced by the *MAT* locus in *Cryptococcus*, including virulence, filamentation, and mitochondrial inheritance during sexual reproduction (38, 67-71). Additionally, the α pheromone genes appear to have a significantly higher intrinsic expression level compared to the **a** pheromone genes in the absence of a compatible mating partner (72), indicating that *MAT*α cells might be better poised to undergo sexual reproduction compared to *MAT***a** cells, which could explain why the *MAT*α allele positively influences hyphal growth.

Regulatory networks controlling cellular morphology can be flexible, involving both genetic and epigenetic factors, and undergoing frequent diversification leading to species-or strain-specific adaptation. In our study, while the nonsense mutation in the *RIC8* gene led to non-yeast form growth under mating-inducing conditions in *C. deneoformans, ric8* mutants of the laboratory *C. neoformans* strains H99 and KN99**a** continue to grow as yeasts under the same conditions. It has also been shown in a previous study that *ric8* deletion compromises titanization in *C. neoformans* (46). Additionally, selfing and unisexual reproduction have yet to be identified in *C. neoformans*, although constitutive activation of the *RAS1* gene, encoding a RAS family GTPase, has been shown to lead to self-filamentous hyphal growth through cAMP, MAP kinase, and RAS-specific signaling cascades (58). Thus, it is possible that other genetic elements present in *C. deneoformans*, but absent in *C. neoformans*, act in concert with the *RIC8* gene to influence the selfing phenotypic variation observed in this study. Such genetic factors would not have been detected if they are fixed in *C. deneoformans* or in both parental strains used to generate the mapping population.

Such gene network plasticity and diversification has also been observed in other species. For example, the standard *Saccharomyces cerevisiae* laboratory strain S288C is defective in yeast-pseudohyphal dimorphism, and studies have shown this defect is due to a nonsense mutation in the *FLO8* gene, which likely occurred during laboratory cultivation. This mutation also blocks flocculation, invasive growth by haploid cells, and pseudohyphal growth in diploids cells (73). Further studies showed the cell-cell and cell-surface adhesion phenotype controlled by the *FLO* gene family are regulated both genetically and epigenetically through a network involving activators that undergo frequent loss-of-function mutations as well as transcription factors whose genetic variation allows diversification of this regulatory circuit among natural isolates (73-75). The four B3502 strains analyzed here are morphologically similar when grown on rich medium under standard conditions, with the only noticeable difference being that colonies from strain B3502_C have a smoother and shinier surface than those from the other B3502 strains. Thus, it is possible that the selection of the spontaneous nonsense mutation in the *RIC8* gene and the nonsynonymous substitution in the gene *CNC06490* were accidental during laboratory passage and strain construction. Nevertheless, our study demonstrates that genetic variation leading to novel phenotypes can evolve rapidly through changes in key regulatory pathways. More importantly, the epistatic interactions between the QTGs and the *MAT* locus identified in our study illustrate the importance of considering background effects when studying genetic variation and phenotypic outcomes. Abundant genetic polymorphisms have been identified in natural populations of every species analyzed, including those in the *Cryptococcus* species complex (24, 76-80), which is accompanied by extensive phenotypic variation among the strains. Interestingly, phenotypic variation in processes including melanin production, stress response, drug resistance, and sexual fertility, have been observed in laboratory evolved populations of *Cryptococcus*, even among strains of a commonly used *C. neoformans* laboratory strain, H99, which have been maintained in different laboratories (65, 78, 81). While the genetic basis of some of this phenotypic variation has been identified, others remain elusive. It is possible that the genetic regulation of this phenotypic variation is complex and involves myriad interactions between genes and regulatory pathways, whose characterization could be challenging for classic genetic approaches that focus on single genes. A future challenge will be to capitalize upon advances in research approaches and analytical tools to dissect these networks that connect genetic variation with resulting phenotypic outcomes.

## Materials and methods Strains and media

All strains analyzed in this study were grown on YPD solid medium at 30°C, unless otherwise stated. Strain CF830 (JEC21, *NOP1-GFP-NAT*) was generated in a previous study and has a *NOP1-GFP-NAT* construct randomly inserted into the JEC21 genetic background (37).

### Mating and spore dissection

Crosses between strains of opposite mating types, selfing assay of the meiotic progeny from the crosses of B3502 strains, as well as dissection of basidium-specific or random basidiospores were carried out as previously described (82). Briefly, 5 µl of cell suspension which contains cells of the two parentals strains or a single strain for crosses and selfing assay, respectively, was spotted onto MS solid medium plate and incubated in the dark. Plates were inspected daily from day 5 to day 21 for signs of sexual reproduction, including hyphae production as well as basidia and basidiospore chains formation.

### *In vitro* and *in vivo* titan cell induction

*In vitro* titanization assay were carried out as described in previous studies (46). For *in vivo* assays, mice were inoculated intranasally with 5×10^5^ cells suspended in 25 µl of PBS. Six mice were included for each strain, with 3 sacrificed at 8 days post infection and 3 sacrificed at 21 days post infection for histopathology analysis of the lungs.

Animal studies were conducted in compliance with guidelines issued by Duke’s Institutional Animal Care and Use Committee (IACUC) and the United States Animal Welfare Act, under protocol A148-19-07 approved by IACUC.

### Genomic DNA purification and genotyping

Genomic DNA extraction, genotyping of the *MAT* locus, as well as genotyping of the *RIC8* locus were carried out as described in previous studies (71, 83, 84). The **a** and α *MAT* alleles were identified using PCR markers, while the wild-type and nonsense mutant alleles of the *RIC8* gene were differentiated using PCR-RFLP markers (Figure S2).

### Sequencing, aligning, and variant calling

In total, 25 progeny from the B3502_A x CF830 cross, 55 progeny from the B3502_B x CF830 cross, the four B3502 stocks, the parental strain CF830, along with strains JEC20, JEC21, NIH12, and NIH433 were sequenced at the Duke University Sequencing Core Facility using an Illumina platform. For each of the above samples, paired-end, 100 base-pair sequenced reads were aligned to an JEC21(α) *C. deneoformans* reference genome (85) using BWA (v0.7.12-r1039, (86)) and binary alignment maps were constructed using SAMtools (v0.1.1996b5f2294a, (87)). Variant calling was carried out using the FreeBayes (v1.2.0, (88)) haplotype caller, resulting in 114,223 unfiltered genetic variants, the majority of which are between the progenitor strains NIH12 and NIH433 (Supplemental Figure S1).

### Genetic variant filtering and marker generation

The 114,223 genetic variants identified between NIH12 and NIH433 were further filtered to identify those segregating within the 80 F_1_ progeny from the B3502_A x CF830 and B3502_B x CF830 crosses. This was done by retaining only those variants that: 1) were segregating within the 80 progeny; 2) were called across all progeny (call rate equal to 100%); 3) had a read depth per variant of greater than or equal to 10x and an allelic read depth ratio of greater than or equal to 90%; and 4) were at least 1 kb distal from the centromeres (89). This filtering criteria identified 1,814 bi-allelic variants segregating within the F_1_ progeny.

For haplotype and genealogy analysis of progenitor strains, the raw 114,223 genetic variants were filtered in the same manner as described above, however, the filtration criteria were only applied to sequencing from the progenitor strains, NIH12 and NIH433, the B3502 stocks, the JEC20 and JEC21 strains, and the parental strain CF830. This resulted in 103,435 bi-allelic variants which represent high-quality variants between NIH12 and NIH433.

For pair-wise analysis of variants between strains and the B3502 stocks, variants were only called between strains in the comparison in addition to passing all the above filtering criteria.

### Progeny filtering and marker generation

Each of the 80 progeny were filtered to remove progeny that were genetically identical (clones) and those whose phenotypes were not consistent across replicates. This analysis identified twelve strains that separate into five groups of clones, seven of these strains were removed from analysis while five unique strains were kept (one from each clonal group). Four progeny were removed from analysis because their phenotypes were not consistent across experiments, varying between growth in the yeast cell form and the transgressive selfing form. Two progeny were identified as having potential aneuploidy (Chromosome 11 of progeny A03 and Chromosome 14 of progeny A05) but were not heterozygotes across these duplicated chromosomes and thus were retained for QTL analysis. After applying these filtering criteria, 70 progeny and the three parental strains – B3502_A, B3502_B, and CF830 – were used as our mapping population in QTL mapping analysis.

### Information-theoretic measures and QTL mapping

For exploring the association between genotype and phenotype in the F_1_ mapping population, a mutual information statistic based on information-theoretic measures of entropy was utilized in QTL mapping. In information theory, entropy is a measure of the heterogeneity of a random variable (54, 90). For a discrete, random variable *G* with values *g* (such as the bi-allelic state at a given loci (coded here as *g* = 0 if the allele is from CF830 or *g* = 1 if inherited from B3502) and a probability mass function *p*(*g*), the entropy *H*(*G*) is defined as:

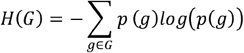

with units “nats” when using this natural log formulation. With two random, discrete variables *G* and *S* (where *S* is a set of discreetly coded phenotypes such as 0, 1, 2, etc.) the joint and conditional entropies are defined (respectively) as follows:

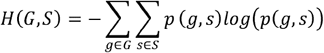

and

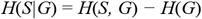

Given these, the mutual information between a genetic variant site, *G* and phenotypic scores, *S* – which measures the reduction in uncertainty of *S* given knowledge about *G* – is calculated as: *I*(*G, S*) = *H*(*S*) *− H*(*S*|*G*) and used here as a measure of the main effect of the genetic variation at variant *G* on the phenotypic classes in *S* (54, 90). This metric was utilized genome-wide for each of the 1,814, bi-allelic variant sites (*G*_*n*_) segregating in the F_1_ progeny. This analysis was conducted using the Python package scikit-learn (91) and repeated for various coding schemes of the selfing and cellular growth phenotypes such as binary (yeast : 0 and non-yeast forms : 1), ternary (yeast: 0, large cell: 1, selfing: 2), or quaternary (yeast: 0, large cell: 1, selfing: 2, transgressive selfing: 3) scores. A 99% confidence interval for QTL on Chromosome 14 was calculated using the binary phenotypes as described in Visscher et al. (92), sampling a thousand times with replacement and taking the location of maximum variant location on that QTL.

### Permutation tests

Permutation tests, as described in Churchill and Doerge (93), were conducted to establish significance thresholds for the mutual information between genotype and the morphological phenotypes (yeast, large cell, selfing and transgressive selfing) from QTL mapping. The number of permutations was ten thousand. The 99th percentile of the permuted distributions of genotype-phenotype associations was used to estimate the thresholds for significance.

### Analyzing three-way epistasis

An information-theoretic approach proposed by Hu et al. (54) was used to examine evidence of three-way genetic epistasis and association of cellular growth phenotypes with genotypic variation at the QTN in *RIC8* and *CNC06490*, as well as the *MAT* locus (*MAT*α = 0, *MAT***a** = 1). Briefly, “information gain” (*IG*) is measured in terms of mutual information and is a measure of the knowledge gained from non-linear, synergistic, epistatic interactions (54). We applied this method using the quaternary discretization of growth phenotypes via the Python package scikit-learn (91). One thousand permutations, which permuted the genotypes of progeny within each phenotypic class, randomizing any non-linear interactions while preserving the main effects, were used to establish significance. From these permutations, the *p*-value for three-way epistasis was calculated as the fraction of permuted *IG* statistics greater than the observed.

## Data availability

Raw sequence reads generated from samples utilized in this study are available on NCBI’s Sequence Read Archive (https://www.ncbi.nlm.nih.gov/sra) under BioProject identification number PRJNA747819. The software developed for both QTL analysis and figure generation are publicly available on GitHub: https://github.com/magwenelab/Self-Filamentation_B3502_progeny.

## Acknowledgements

The authors thank Drs. Sue Jinks-Robertson, Thomas Petes, Andy Alspaugh and Vikas Yadav and Shelby Priest for their critical reading and comments on the manuscript. We are grateful to Drs. Elizabeth Ballou and Ci Fu for their generous assistance and suggestions on *in vitro* titanization assays and FACS analyses, respectively. This study was supported by NIH/NIAID R01 grants AI39115-24 and AI50113-16 awarded to JH and R01 grant AI33654-05 awarded to JH and PM. JH is Co-Director and Fellow of the CIFAR program Fungal Kingdom: Threats & Opportunities.

## Supplemental Figure Legends

**Supplemental Figure S1. Genetic variants identified among the B3502 stocks**.

(A) *C. deneoformans* progenitor strain haplotypes. Between the strains NIH12 (blue) and NIH433 (orange), 104,153 genetic variants were identified and used to construct haplotypes per chromosome (x-axis) for the B3502 stocks, the congenic strains JEC20 and JEC21, and the transformed strain CF830 (y-axis, top to bottom, respectively). The location of the *MAT* locus is annotated in red and its approximate boundaries on Chromosome 4 are defined by a red horizontal bar. The two black arrows at the bottom of Chromosome 2 indicate the genomic regions in which the vast majority of the polymorphisms between JEC21 and B3502 were identified. (B) Shown here are genetic variants identified in pairwise comparison between B3502 stocks and the JEC21 reference genome. The centromeric regions were excluded due to the presence of repetitive elements and transposable elements for which variants are difficult to be called with high confidence using short-read sequences. The *MAT* locus was also excluded in comparisons between B3502 stocks and the JEC21 reference genome, given the highly diverged *MAT***a** and *MAT*α alleles in B3502 and JEC21, respectively.

**Supplemental Figure S2. Segregating genetic variation in B3502 stocks and F**_**1**_ **progeny**. Genotypes of genetic variants (x-axis) segregating within F_1_ progeny from crosses between B3502_A x CF830 and B3502_B x CF830 are depicted. Genotypes for strains (y-axis) are colored blue if they are identical to the *C. deneoformans* strain NHI12, and yellow if they have the alternative alleles. The haplotypes of the B3502 stocks, the parental strains CF830, reference strain JEC21, and ten of the F_1_ progeny are shown.

**Supplemental Figure S3. Markers used to genotype the *MAT* and *RIC8* loci**.

Shown here are markers used to genotype the *MAT* locus and the *RIC8* locus. On the left are genotyping results of the B3502 stocks and the JEC20 and JEC21 control strains. On the right are genotyping examples of 24 meiotic progeny dissected from B3502_A x CF830 crosses. The **a** and α alleles of the *MAT* locus are differentiated by two PCR markers targeting the **a** and α alleles, respectively; the *RIC8* wild-type and nonsense mutant alleles are differentiated by a PCR-RFLP marker (digestion with restriction enzyme Hpy188I).

**Supplemental Figure S4. Diameters of cells from different strains**.

For each strain, on the right is a scatter plot showing the distribution of cell sizes when grown under *in vitro* titan cell inducing conditions, and on the left is a box-and-whisker plot of the same dataset, with the box representing data between the 25% and 75% percentiles, and the line and cross within the box representing the median and mean of the population, respectively. More than 140 cells from each strain were analyzed. Significance was assessed using Mann-Whitney U Test.

**Supplemental Figure S5. Manhattan plots of genome-wide strength in association between genetic variants and progeny phenotypes**.

(A) Distributions of binary (yeast and non-yeast growth), ternary (yeast, large cell, and selfing), and quaternary (yeast, large cell, selfing, and transgressive selfing) phenotypes used in QTL mapping. (B) Results of QTL analysis using mutual information statistic (y-axis) to associate variation at genetic variant sites (x-axis) and binary, ternary, and quaternary phenotypes. Colors separate chromosomes. The approximate location of the *MAT* locus (on Chromosome 4) is marked by a horizontal red line.

**Supplemental Figure S6. Gene model of *CNN00400***.

From QTL analysis a SNP within the gene *CNN00400* at 1,740 bp was identified and is predicted to cause a nonsynonymous change – glycine to arginine – near the end of the first exon.

**Supplemental Table S1. Genetic variants identified in the genomes of B3502 stocks**

**Supplemental Table S2. Analyses of the progeny from crosses B3502_A** × **CF830**

**Supplemental Table S3. Analyses of the progeny from crosses B3502_B** × **CF830**

**Supplemental Table S4. Analyses of the progeny from crosses B3502_C** × **CF830**

**Supplemental Table S5. Analyses of the progeny from crosses B3502_D** × **CF830**

