## Supplementary figures and images for "Epistatic genetic interactions govern morphogenesis during sexual reproduction and infection in a global human fungal pathogen"

### Supplemental Figure S1

#### Suppl. Figure S1

**A**

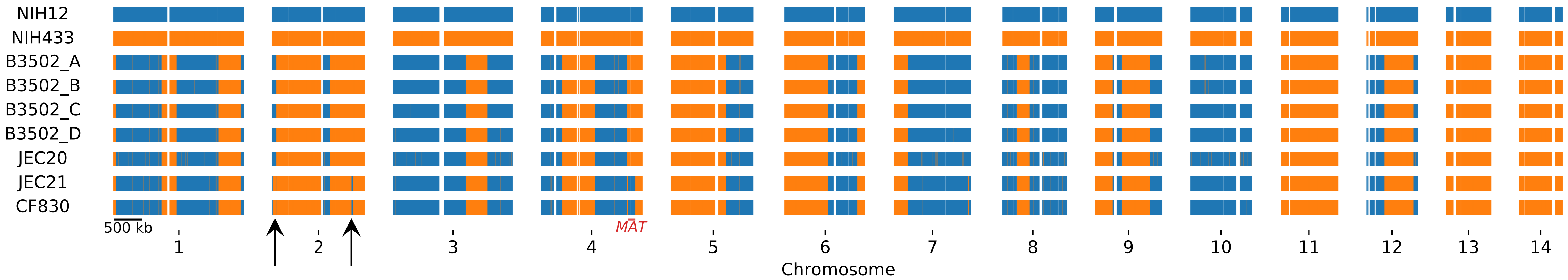


**B**

| | CF830 | B3502_A | B3502_B | B3502_C | B3502_D |
|---|---|---|---|---|---|
| **JEC21** | 2 | 508 | 533 | 505 | 502 |
| **CF830** | | 510 | 535 | 508 | 504 |
| **B3502_A** | | | 5 | 19 | 25 |
| **B3502_B** | | | | 23 | 29 |
| **B3502_C** | | | | | 14 |

### Supplemental Figure S2

#### Suppl. Figure S2

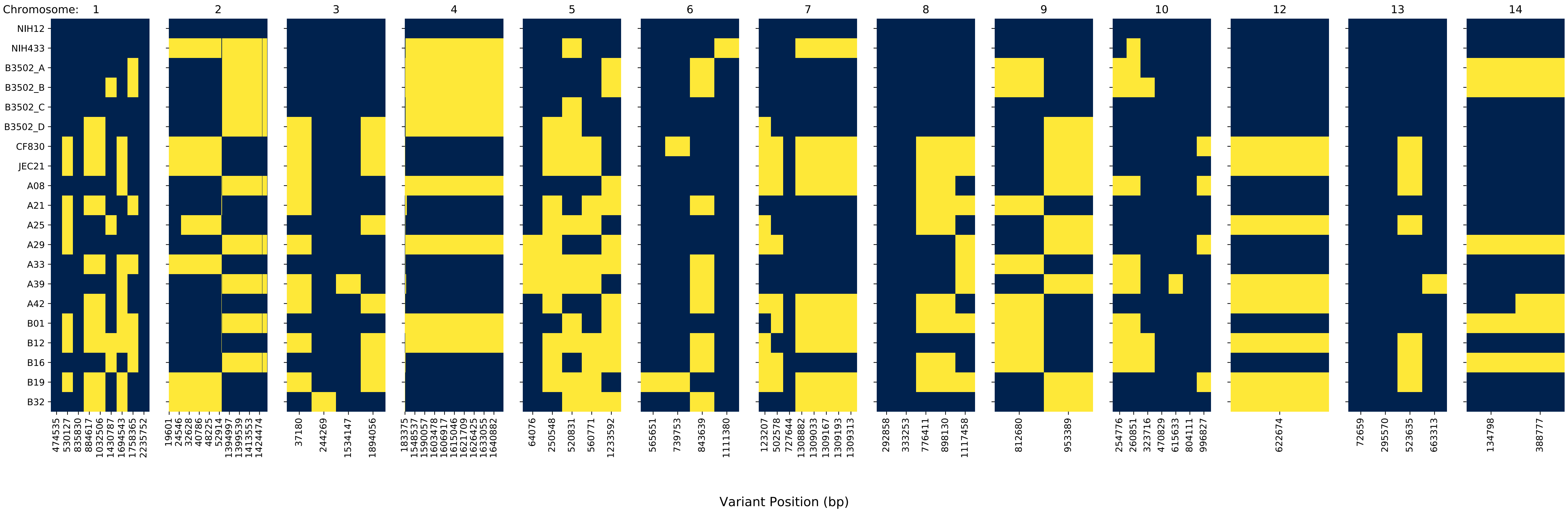

### Supplemental Figure S3

Suppl. Figure S3

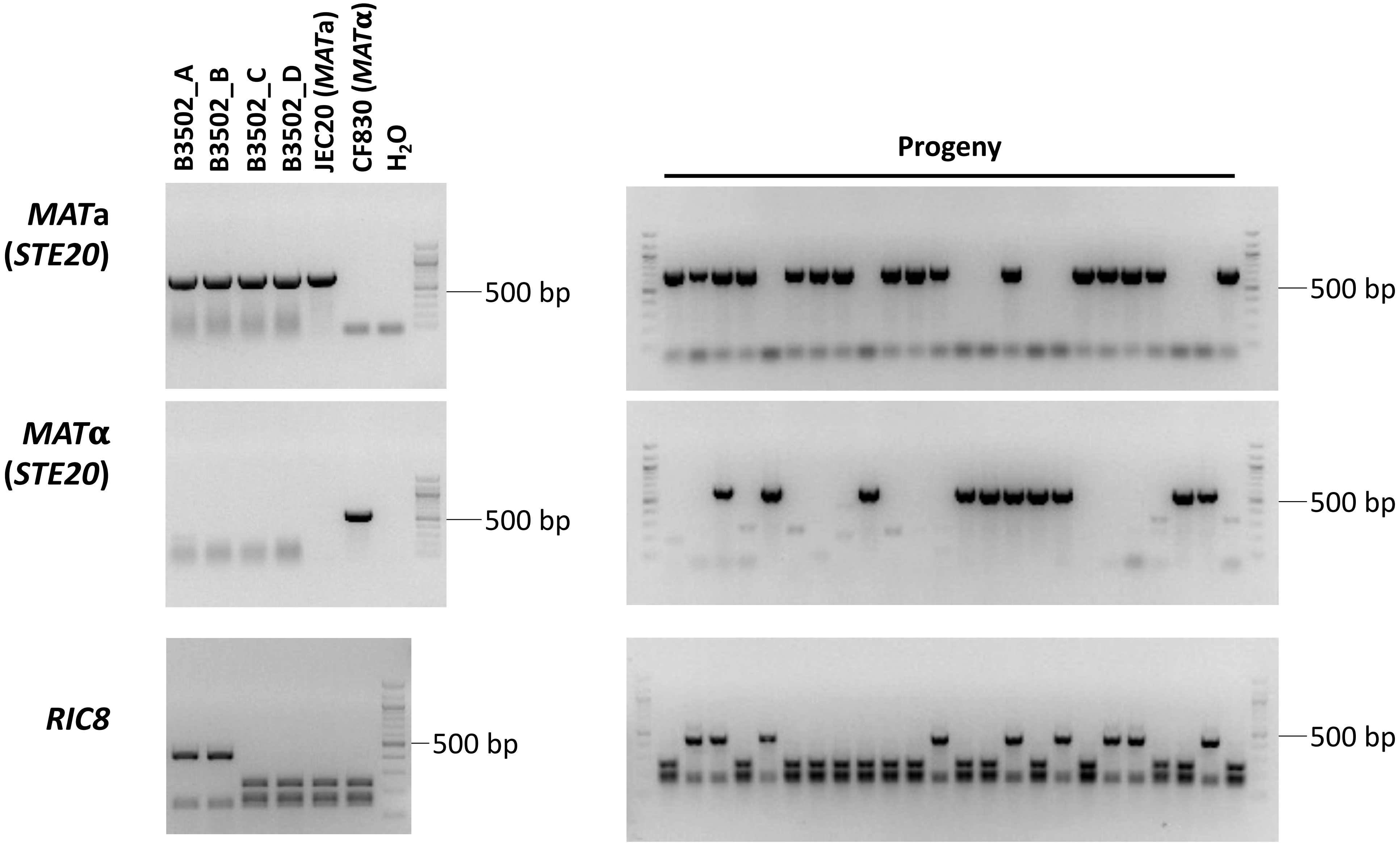

### Supplemental Figure S4

Suppl. Figure S4

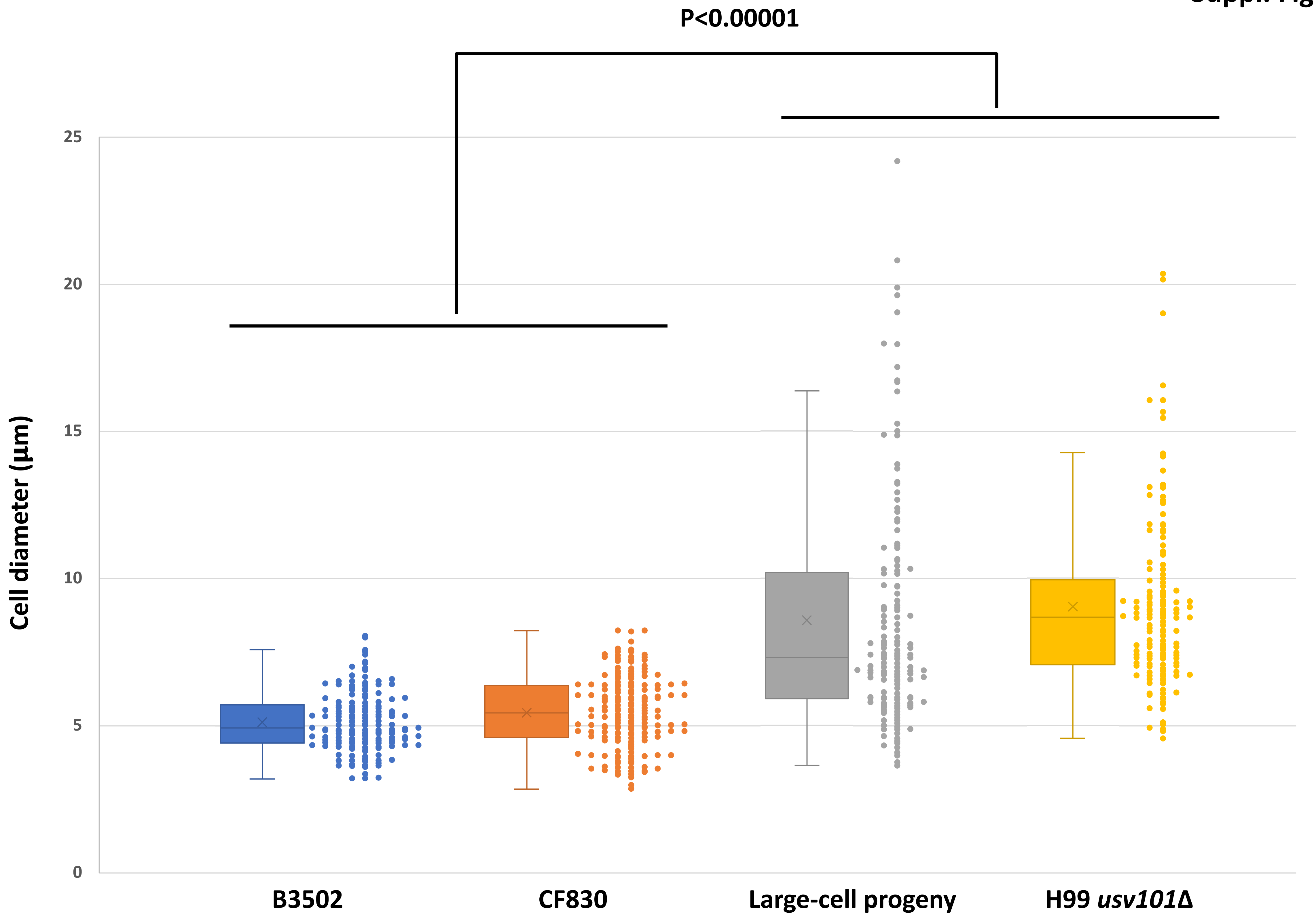

### Supplemental Figure S5

#### Suppl. Figure S5

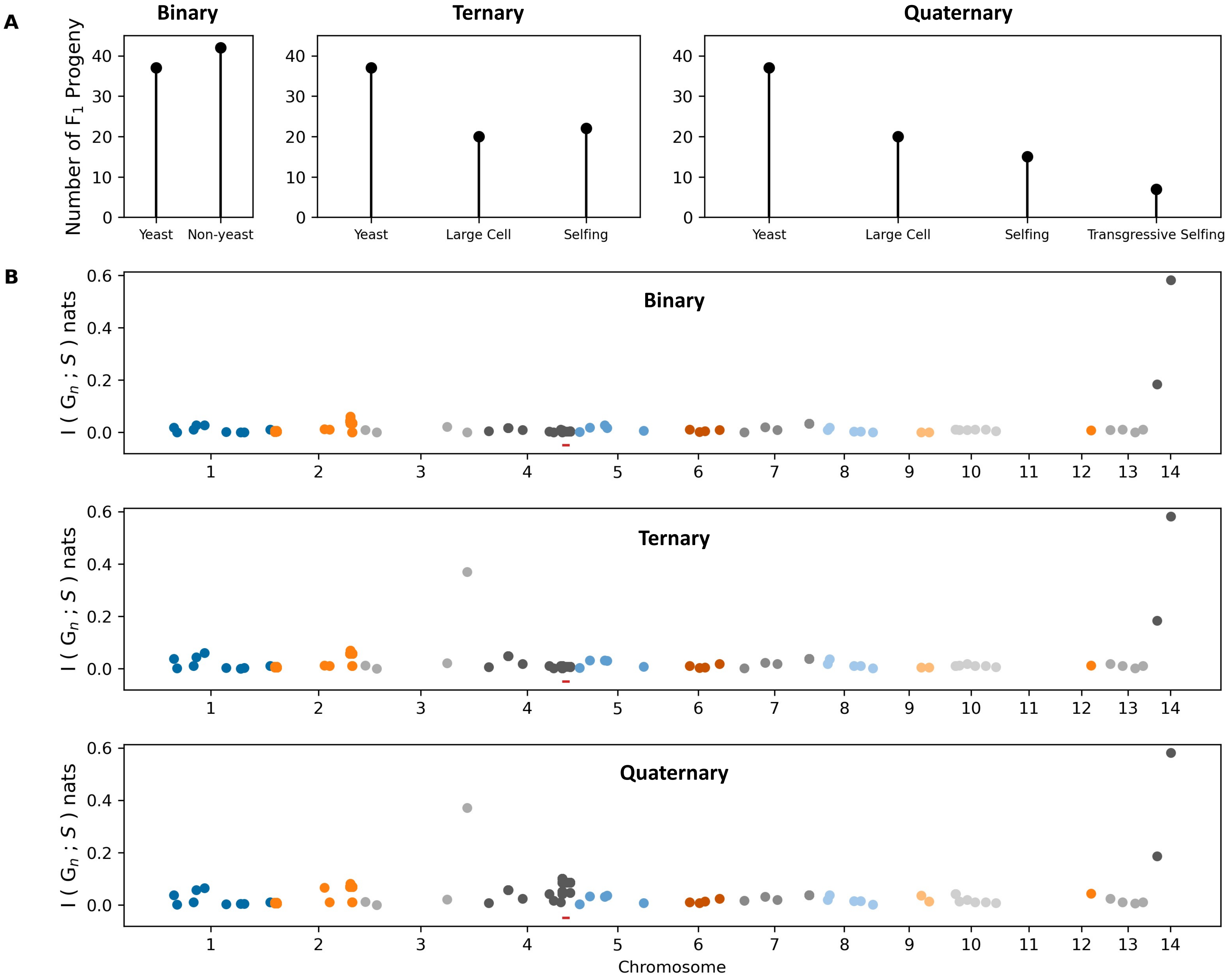

### Supplemental Figure S6

**Suppl. Figure S6**

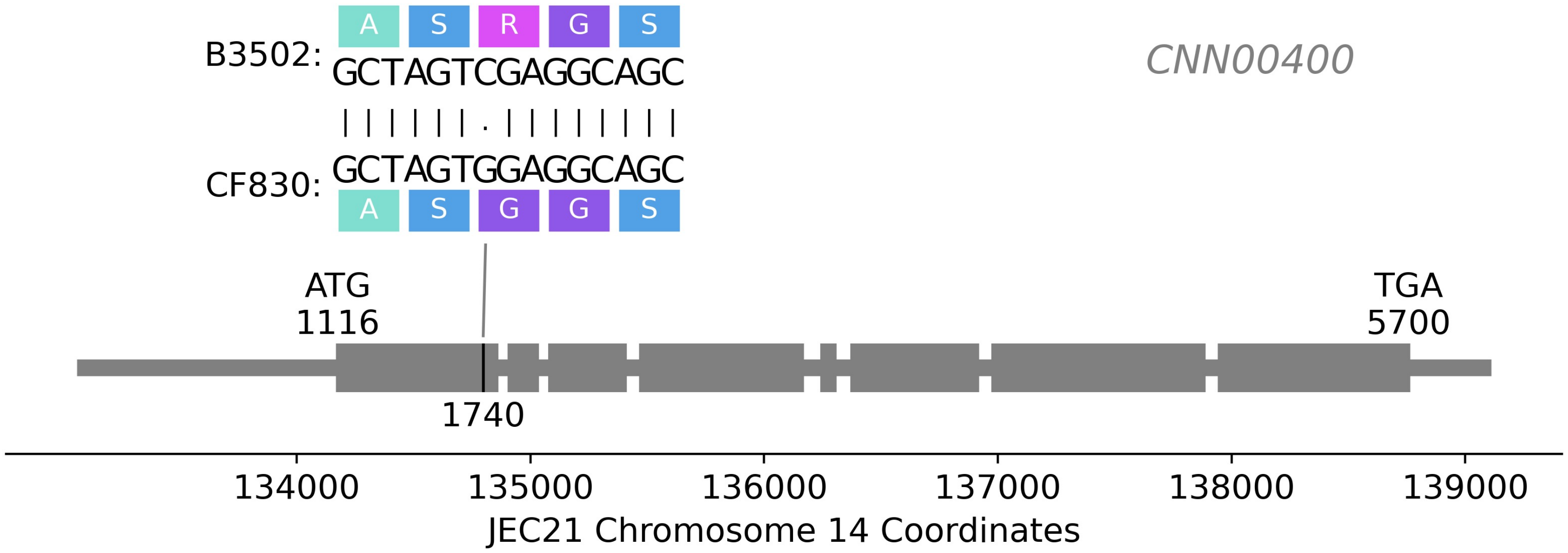
